# Snowmobile noise alters bird vocalization patterns during winter and pre-breeding season

**DOI:** 10.1101/2023.07.13.548680

**Authors:** Benjamin Cretois, Ian Avery Bick, Cathleen Balantic, Femke B. Gelderblom, Diego Pávon-Jordán, Julia Wiel, Sarab S. Sethi, Davyd H. Betchkal, Ben Banet, Tor Arne Reinen

## Abstract

Noise pollution poses a significant threat to ecosystems worldwide, disrupting animal communication and causing cascading effects on biodiversity. In this study, we focus on the impact of snowmobile noise on avian vocalizations during the non-breeding winter season, a less-studied area in soundscape ecology. We developed a pipeline relying on deep learning methods to detect snowmobile noise and applied it to a large acoustic monitoring dataset collected in Yellowstone National Park. Our results demonstrate the effectiveness of the snowmobile detection model in identifying snowmobile noise and reveal an association between snowmobile passage and changes in avian vocalization patterns. Snowmobile noise led to a decrease in the frequency of bird vocalizations during mornings and evenings, potentially affecting winter and pre-breeding behaviors such as foraging, predator avoidance and successfully finding a mate. However, we observed a recovery in avian vocalizations after detection of snowmobiles during mornings and afternoons, indicating some resilience to sporadic noise events. These findings emphasize the need to consider noise impacts in the non-breeding season and provide valuable insights for natural resource managers to minimize disturbance and protect critical avian habitats. The deep learning approach presented in this study offers an efficient and accurate means of analyzing large-scale acoustic monitoring data and contributes to a comprehensive understanding of the cumulative impacts of multiple stressors on avian communities.

**Graphical Abstract:** 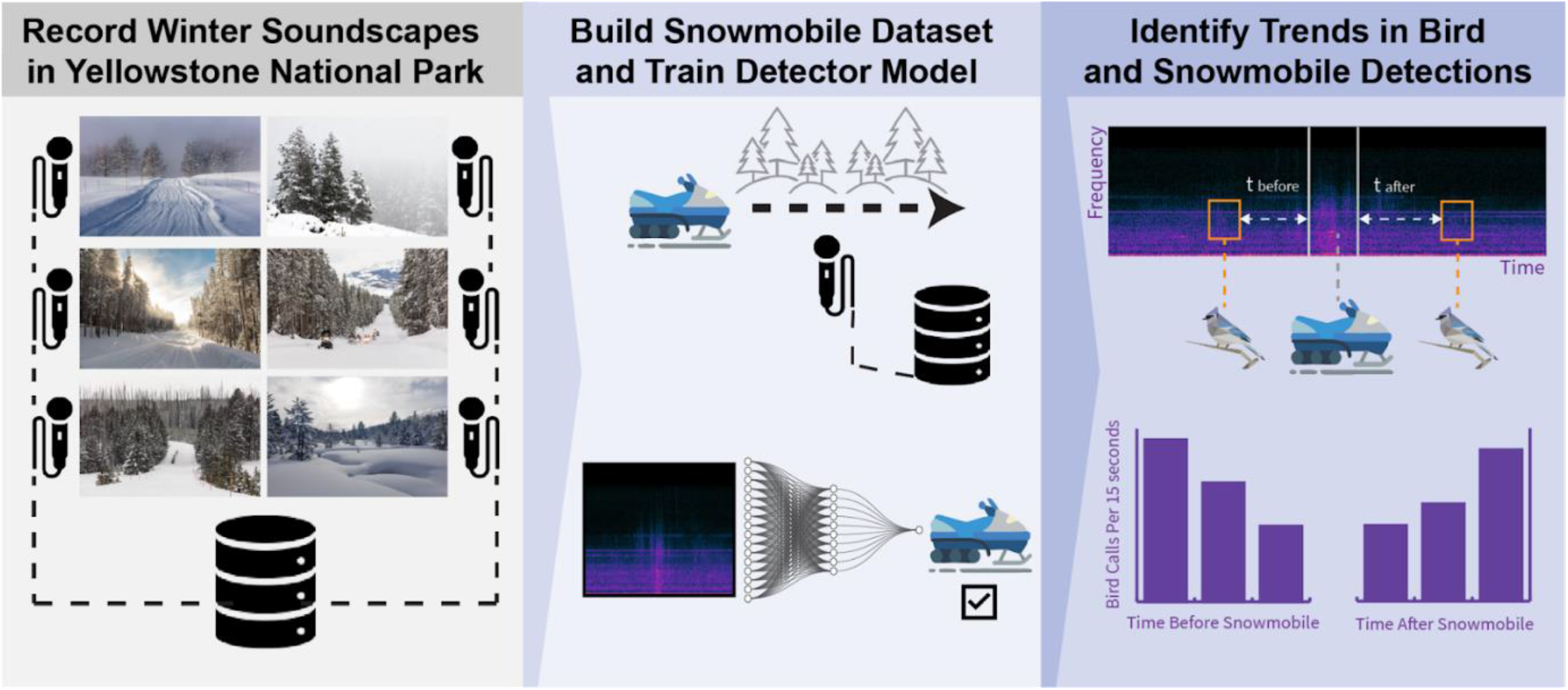

## 1. Introduction

Noise pollution is a growing concern for ecosystems worldwide. Anthropogenic noise from sources such as traffic can mask biophony, disrupting animal communication and causing cascading effects on habitat availability (Bayne et al., 2008; Ware et al., 2015), reproductive success (Naguib, 2013; Schroeder et al., 2012) and biodiversity (Ware et al., 2015). Passive acoustic monitoring methods are uniquely suited to study such interplay between biophony and anthrophony, as they enable cost-effective collection of soundscape data over long periods of time and wide geographic areas.

However, this wealth of data creates a need for automated techniques to identify and separate target sounds such as vocalizations from species of interest or anthropogenic noises. Deep learning methods are well suited for this task and have been shown to successfully identify vocalizing animal species (Kahl et al., 2021; Poupard et al., 2022; Stowell et al., 2019), human voices (Cretois et al., 2022), and engine noises (Espinosa et al., 2021) from audio recordings. By using multiple automated detectors in parallel, one may efficiently detect and study relationships between anthropogenic sounds and animal vocalizations.

Previous acoustic ecology studies have demonstrated the complex effects of anthropogenic noise on bird species: reduced body condition (Ware et al., 2015), increased vigilance and lower food intake (L. Quinn et al., 2006), increased call duration (Courter et al., 2020), reduced nest visitation (Naguib, 2013), and habitat quality reduction (Ware et al., 2015). Most studies have focused on chronic effects of noise which have persisted for weeks, months, or years. Previous studies that have explored immediate effects of noise on wildlife, such as for snowmobile and aircraft noise events (Borkowski et al., 2006; Lawler et al., 2004; Maier et al., 1998; Vincelette et al., 2021), have relied on direct observation or manual sound annotation, requiring many work hours and potentially limiting temporal scope. Deep learning techniques have made it possible to analyze large amounts of data for event identification and classification (Stowell, 2022). This presents an opportunity to study the impact of anthropogenic noise on avian vocalizations at both a higher temporal resolution and with a larger temporal and geographic scope.

Most literature on noise impacts to birds specifically considers breeding season vocalizations (Oden et al., 2020; Shannon et al., 2016), while impacts of noise during the non-breeding winter and pre-breeding seasons remain understudied (though see Mullet et al., 2016; Quinn et al., 2022) despite the non-breeding season comprising a substantial portion of avian life history (Faaborg et al., 2010). Because noise and more specifically anthropogenic noise such as engine noises may act as a threat multiplier in combination with other stressors like habitat loss and climate change, it is critical to better understand the extent of the influence of anthropogenic winter noise on avian vocalizations. Automated detection models for anthropogenic winter noise (such as engine noise from snowmobiles) can facilitate efficient assessments of avian responses to noise and inform natural resource management strategies.

Reliable automated detection of snowmobiles in particular presents unique challenges due to both a lack of available training data for deep learning models and confounding noise sources in the soundscape. Wind noise is a particular issue, as it introduces low-frequency acoustic energy that may be mislabeled as an engine (Juodakis & Marsland, 2021). While studies have mapped the prevalence of snowmobile noise in American wilderness areas (Mullet & Morton, 2021) and explored potential for masking of bird calls (Keyel et al., 2018), little work has been done to measure direct impacts of snowmobiles on bird vocalizations (National Park Service, 2013).

Yellowstone National Park, located in the western United States, is a unique place spanning 8900 km^2^ where wildlife coexists with anthropogenic noise, including along roads. Snowmobiling is a popular recreational activity in the park, attracting thousands of visitors annually. Between mid-December and mid-March, the park’s roads are only accessible via such “oversnow” vehicles such as snowmobiles and snowcoaches. Snowmobiles often generate more noise than car engines (Daily & Raap, 2002; Menge et al., 2002) and have been shown to instigate vigilance, travel, and occasional flight responses in Yellowstone’s elk and bison, especially when animals were closer to roads (Borkowski et al., 2006). Noise associated with oversnow vehicles is an important management concern in the park and noise impacts have been routinely measured via acoustic monitoring (Burson, 2018). To manage and mitigate oversnow vehicle impacts, Yellowstone implemented a series of environmental impact statements and environmental assessments, culminating in the adoption of the current Winter Use Plan in 2013 (National Park Service, 2013). However, it is unclear how snowmobile noise impacts the park’s diverse winter avifauna such as owls, raptors, and overwintering songbirds such as bohemian waxwings, dark-eyed juncos, red-breasted nuthatches, and mountain chickadees (National Audubon Society, 2020; National Park Service, 2013; Walker et al., 2020).

In landscapes where recreation and wildlife conservation may present conflicting priorities and legal challenges (Dustin & Schneider, 2004), a rigorous methodology is necessary to measure and assess noise impacts. Acoustic monitoring of winter soundscapes provides one avenue for evaluating noise impacts, but it can be challenging to efficiently identify target noise and sound events in large audio datasets. Addressing these obstacles may offer novel insights on the impacts of snowmobile noise on non-breeding birds and support natural resource managers in making informed decisions that balance recreation and conservation.

In this study, we developed a methodology for assessing noise impacts from snowmobiles on the winter and pre-breeding avian community. We used acoustic monitoring data to pursue the following objectives: (1) train a deep learning model to detect snowmobile noise, (2) test the snowmobile detector on a large winter acoustic monitoring dataset collected at Yellowstone National Park (Wyoming, USA), and (3) assess the methodology for investigating how bird vocalization frequency reacts to and recovers from snowmobile noise, using the Yellowstone dataset as a case study.

## 2. Material and Methods

**Fig 1.**
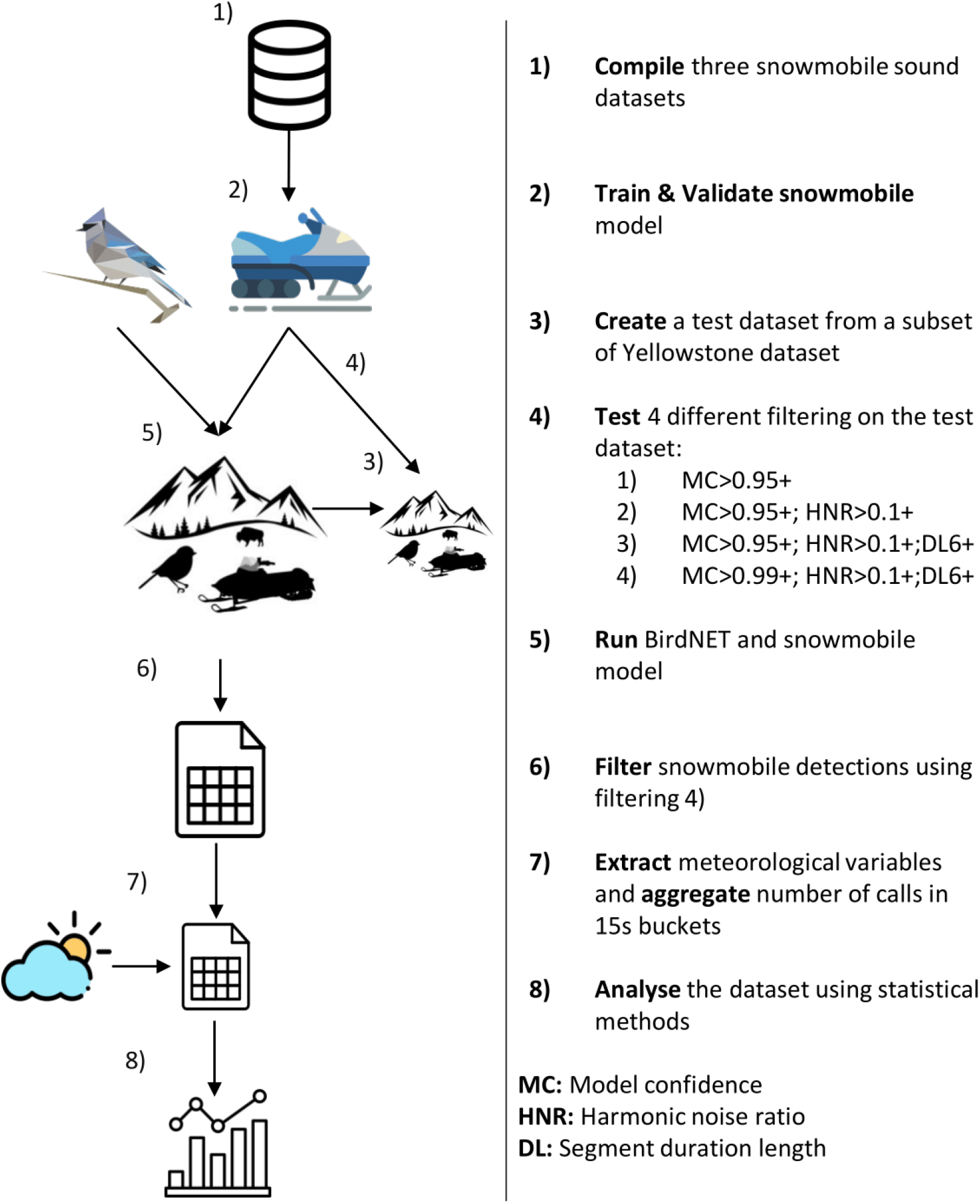
Workflow of the method.

### 2.1. Data Collection

#### 2.1.1 Dataset 1: Training and Validation of the Model

To obtain data for training and validation, we compiled three audio datasets (Fig 1-1). 1) During the winter of 2022 in Bymarka near Trondheim, Norway, an Audiomoth (model 1.1.0) was placed 1.20 m above the ground in the middle of a 150×150 meters skiing play area for children. A single 4-stroke snowmobile (Lynx Yeti ACE 900) transited the area at an approximate speed of 10 km/h. The audio also contains singing birds, people talking, and children playing. 2) The data from Hattfjelldal commune in Nordland, Norway, were recorded with an Audiomoth 1.1.0 in March and April 2022 at four sites along snowmobile tracks. Snowmobile traffic was recorded for a few hours at each site. 3) A third dataset from previous work gathered audio recordings of passing snowmobiles with different speeds and engine types (Gjestland & Haukland, 2017). All files were analyzed with Raven Pro 1.6. Some were discarded because of bad quality or clipping.

The recordings contained both segments with snowmobile and without snowmobile passages and were therefore manually reviewed and labeled. All time intervals where a snowmobile was audible were categorized as “snowmobile”. All time intervals where no snowmobile was audible were categorized as “background”. In total, the training and validation sets were composed of 4 and 33 hours of snowmobile sounds and background sounds, respectively.

#### 2.1.2. Dataset 2: Model Testing and Use in a Passive Acoustic Monitoring Scenario

To evaluate the performance of our classifier we used a large acoustic dataset composed of continuous recordings (i.e. no time gaps between files) collected at 20 sites in Yellowstone National Park. Data collection sites occurred in the vicinity of two roadside locations (Madison Junction and near Lewis Lake) and one destination area (Old Faithful) (See supplementary map 1). Recordings occurred during the winter use season spanning from December 15th until March 15th and between the years 2008 and 2017. Because of the variation in sampling protocol at each site, there are inter-site differences in the time range (i.e. some sites containing acoustic data spanning from 2009 up to 2017 while other sites containing only a year of acoustic data) and in the length of the recorded files (i.e. some files containing about 15 seconds of audio while others containing about 12 hours of audio) (Burson, 2009, 2018). In total the Yellowstone dataset comprised approximately 43,000 hours (equivalent to 1791 days) of audio collected through passive acoustic monitoring. For simplicity we refer to this dataset as the Yellowstone dataset hereafter.

The test dataset was composed of 36 hours of randomly sampled data from the Yellowstone dataset that we manually labeled using Raven Pro (Fig 1-3). All time intervals where a snowmobile was audible have been noted. In addition, if other audible engine sounds such as helicopter noise, aircraft noise, and car noise were detected, they were labeled as well. Using the model on the test dataset allowed us to infer the model’s performance metrics including precision, recall and F1 score.

The portion of the Yellowstone dataset not included in model testing was used to study the impact of snowmobiles on avian vocalization. No manual tagging was available and it was unknown whether the audio files contained snowmobile noises or bird vocalization.

### 2.2 Snowmobile Detection Model Architecture and Training

To construct a model efficient at classifying snowmobile noises we used a ResNext model (i.e. Residual network that leverages attention layers) that had been pre-trained on AudioSet as feature extractor (Guzhov et al., 2021). We appended a final fully connected layer with two output neurons. Based on a hyperparameter search using Ray.tune (Liaw et al., 2018), the base ResNext model was then fine-tuned on our training data using the cross entropy loss function, a learning rate of 0.005, and a batch size of 32. The model was trained for a total of 21 epochs as we set an early stopping on the validation loss (i.e. if the validation loss increases after reaching a minimum the model stops training) (Fig 1-2).

The training data was resampled at 44.1 kHz, split into non overlapping 3 second segments and transformed into a log-power spectrogram using a Fast Fourier Transform window size of 46 ms (2048 samples at 44.1 kHz) and a hop length of 13 ms (561 samples at 44.1 kHz). The spectrogram was split into 3 frequency bands: lower (0.00−7.35 kHz), middle (7.35−14.70 kHz) and upper (14.7− 22.05 kHz). The resulting spectrogram dimensions were [3, 341, 234].

Data augmentation is a practice which consists of artificially increasing the diversity of the dataset by slightly modifying the input data used to train the neural network. Data augmentation is especially useful when a low amount of training data is available (Salamon & Bello, 2017). To take advantage of our limited training dataset we made extensive use of data augmentation by using Python’s audiomentations library (Jordal et al., 2022). In particular, we used time and frequency masking (i.e. this acts as a form of dropout of frequencies and time regions), we added time shifting, we applied a low pass-like filterbank with variable octave attenuation (i.e. to simulate the effect of distance), modified the frequency equalization (i.e. to simulate the change in the response characteristics of the microphone) and finally added Gaussian noise in the range [0.001, 0.015]. The probability for each of the augmentation types was set to 0.5.

Outdoor recordings are obtained in uncontrolled environments where extraneous noise signals are bound to contaminate the data. One particularly challenging type of contamination is wind noise, which includes both the acoustic sources that it creates (like the rustling of leaves), and pressure changes at the microphone’s membrane that leave audible artifacts in the recording (Cook et al., 2021). Residual environmental noise such as wind and rain have been a major obstacle for bioacoustic detectors incorporating deep learning models thus far (though see Juodakis & Marsland, 2021; Metcalf et al., 2020) and failure to correctly account for weather noise can dramatically affect detector performance. Various approaches to environmental noise suppression have been developed for different tasks but classic denoising methods such as the Wiener or MMSE filters are not applicable to wind because of its rapid dynamics (Juodakis & Marsland, 2021).

The cyclic nature of a snowmobile’s combustion engine (the fact that it operates at an approximately constant rpm given a short enough time segment), ensures that snowmobile engines produce a characteristic periodic signal, in clear contrast to the nonperiodic wind noise. Periodic signals (i.e. harmonic sounds) can be distinguished from nonperiodic signals (i.e. inharmonic sounds) with harmonicity descriptors. One such descriptor is the harmonic noise ratio (HNR) (Moreau et al., 2006).

Therefore, we additionally computed the harmonic ratio for the lower frequencies (obtained by applying a low-pass butterworth filter) of the recordings. In our case, a lower HNR indicates that the source is nonperiodic, and is therefore more likely to contain wind instead of engine noise. Based on the results from the test dataset we classified a segment as containing snowmobile noise if the confidence of the snowmobile model was greater than or equal to 0.95, and the HNR larger than 0.1.

The pipeline for training the snowmobile detector was made in Python using PyTorch. The model was trained on a NVIDIA A40-24Q GPU.

### 2.3. Analysis of the Yellowstone dataset

#### 2.3.1. Extracting Snowmobile and Bird Detections

To be processed by the snowmobile model, each audio file is first split into non-overlapping three second segments. Each audio chunk is then transformed into a spectrogram using a Fast Fourier Transform window size of 46 ms samples and a hop length of 13 ms. Finally, the spectrogram passes through the snowmobile model which outputs the model confidence of whether a snowmobile was present and the harmonic to noise ratio. In contrast, BirdNET accepts mel-spectrograms as input and each audio file is re-sampled to 48 kHz, split into non overlapping three second segments and converted to a mel scale spectrogram of 64 bands that is passed through the model (Kahl et al., 2021).

It is possible to get a higher precision (i.e. lower rate of false positives) at the expense of a higher recall (i.e. lower rate of false negatives) by increasing the thresholds for both model confidence and harmonic ratio. For our analysis we chose to lower the number of false positives by filtering for detections with high model confidence and high harmonic ratio. To select the model confidence and harmonic ratio threshold, we computed the F1 score for four increasing levels of filtering and chose the highest F1 score: 1) retaining the detections with model confidence > 0.95 (MC 0.95+), 2) additionally retaining detections with harmonic noise ratio > 0.1 (MC 0.95+; HNR 0.1+), 3) additionally retaining detections with length >= 6 seconds (MC 0.95+; HNR 0.1+; DL 6+) and 4) use model confidence > 0.99 while keeping HNR > 0.1 and duration >= 6 seconds (MC 0.9+; HNR 0.1+; DL 6+) (Fig 1-4).

We ran both our snowmobile detector model using the filtering that lead to the best F1-score (i.e. MC 0.9+; HNR 0.1+; DL 6+) and BirdNET Analyzer V2.1, a deep learning model for bird classification in acoustic data (Kahl et al., 2021)(Fig 1-5), on the Yellowstone dataset. BirdNET Analyzer V2.1 was run with default parameters and a custom list of avian species found in Yellowstone (extracted from https://irma.nps.gov/NPSpecies/Search/SpeciesList/YELL, Supporting document 1).

#### 2.3.2. Estimating Trends in Bird Vocalization frequency with regards to snowmobile detections

To estimate the impact of detected snowmobiles on bird vocalizations, we first subsetted all detection data between December 15th and March 15th, the typical oversnow vehicle season at Yellowstone. To eliminate the possibility of false positive BirdNET detections triggered by a snowmobile event, any bird calls occuring during a snowmobile detection at the same site were removed from the analysis. We retained all BirdNET detections that did not occur during snowmobile events and assumed that any detection bias in the BirdNET model would be consistent prior to and after a snowmobile event. For each bird call, we identified the nearest snowmobile detection at that site before and after the call and extracted hourly wind speed, relative humidity and temperature from the Yellowstone Lake weather station (NOAA station P90, Herzmann, 2023, accessed via Iowa State University’s Mesonet system)(Fig 1-7). We aggregated the detections into buckets of 15 seconds by summing the number of detections within each bucket (Fig 1-7). To reflect the phenology of bird vocalizations we added a variable time_of_day of value “AM” (i.e. morning), “MM” (i.e. afternoon) and “PM” (i.e. evening) if the 15s-bucket was located in the interval [6AM to 12PM], [12PM, 4PM] and [4PM, 12AM] respectively. Early morning hours [12AM to 6AM] were not analyzed due to minimal bird vocalization activity and park restrictions on snowmobile use during these hours. We analyzed only the four sites that contained the most detections (i.e. YELLOFWS, YELLMJ23, YELLFOPP and YELLGVLL) (see Supplementary Map 1) as including sites with few detections would unnecessarily increase model uncertainty.

In the absence of concrete studies determining the precise distance at which birds can perceive engine noise, we opted to include bird vocalizations in the ranges [240, 0] and [0,240s] (Fig 2) before and after the detection of a snowmobile resulting in a dataset containing 34941 individual bird calls. This decision is based on extrapolations from human auditory capabilities.

**Fig 2.**
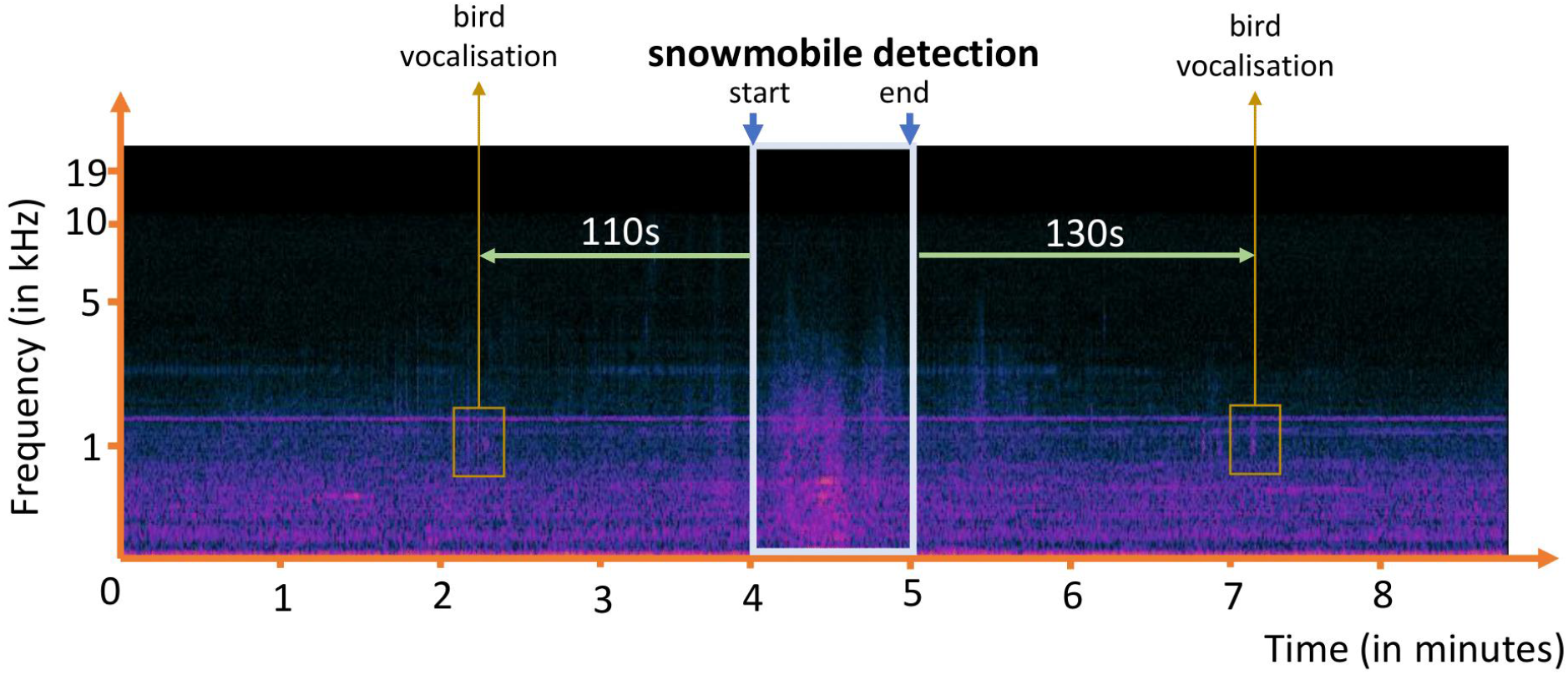
Example of a spectrogram showing the obtention of the distance from bird vocalization (orange square, obtained by BirdNET) to the snowmobile detection (light blue square). The distance from the bird vocalization before and after (green arrow) the snowmobile detection (light blue square) is computed from the start and end of the snowmobile detection.

In ideal conditions such as the relatively quiet environment of Yellowstone National Park, low wind speed and assuming a sufficiently loud snowmobile, research suggests that a human could potentially detect the sound from approximately 2 kilometers away (engine noise can become a nuisance or safety concern at 400-800 meters, suggesting that it’s audible at least this far way, Franks et al., 1996; Miedema & Oudshoorn, 2001). Taking into account a snowmobile traveling at an average speed of 8.33 m/s (i.e., 30 km/h), this translates to humans possibly beginning to hear the snowmobile around 4 minutes (i.e., 2000m / 8.33m/s = 240 seconds) before it passes their location. We used linear models (Fig 1-8) with the log of the number of bird calls (i.e. the number of calls in the 15s bucket) as the response variable (Gaussian error distribution) and the interaction between time before or after detection of a snowmobile and time of day. We chose a Gaussian model over a Poisson model (typical for count data such as ours) because the later model showed a high degree of overdispersion. Log-transforming our response variable solved the problem. To control for the effect of environmental variables on the log-transformed number of bird calls, we included the following variables: hourly wind speed, relative humidity and temperature as explanatory variables. To account for the dependency of observations from the same site and from the same day, we included ‘site’ and ‘date’ as crossed random effects. The nearest hourly weather reading was related to each snowmobile and each individual bird call detection. Any gaps in available weather data were filled using linear interpolation. Sites and dates were included as random effects. All numerical variables were z-score normalized.

The models were fitted using the glmmTMB package (Magnusson et al., 2017) in R 4.2.2 (R Core Team, 2022) and we assessed the fit of the model using the DHARMa package (Hartig & Hartig, 2017).

## 3. Results

### 3.1. Snowmobile Detection Model Performance

Our results show that accounting for wind using a filter on the harmonic ratio value greatly increases model performance (Fig 3). When detections are only filtered using the snowmobile model confidence (>0.95) we obtain a precision of 0.11 (i.e. on average, among 100 positive identifications only 11 are really snowmobiles). However, when filtering detections based on both model confidence (>0.95) and harmonic ratio (>0.100), the precision reaches 0.5. For both MC 0.95+ and MC 0.95+ HNR 0.1+, the recall is of 0.98 (i.e. on average, the model correctly detects 0.98% of all snowmobiles). Accounting for detection length (detections that are at least 6 seconds long) in addition to model confidence and harmonic noise ratio further increases the precision while slightly decreasing the recall (precision = 0.65; recall = 0.94). The final set of parameters filtered by model confidence >0.99 and yielded an increased precision (0.78) while reducing the recall (0.77), displaying the trade off between these metrics.

**Fig 3:**
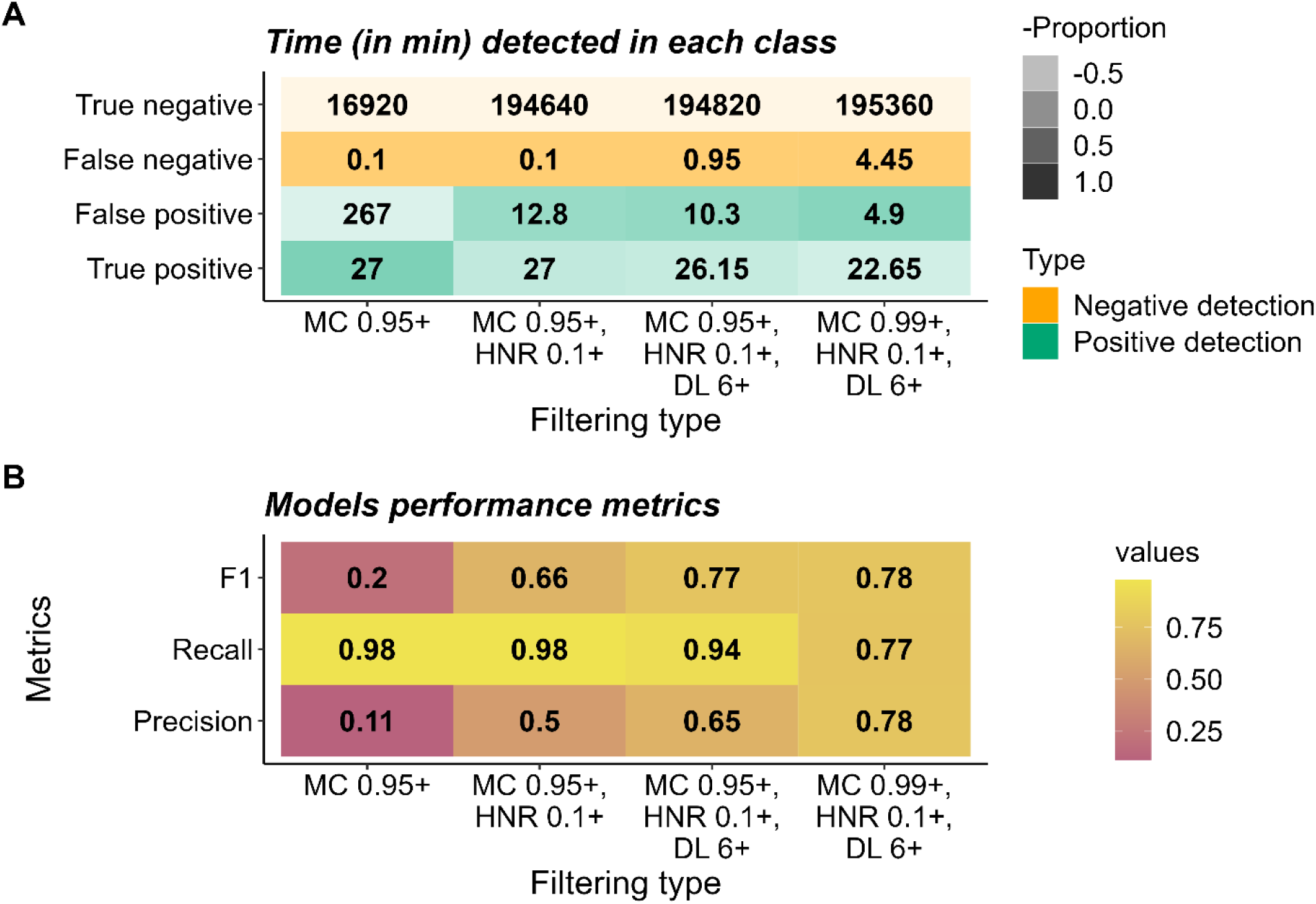
Confusion matrix (A) for snowmobile event detection on the test dataset after different levels of filtering and (B) figure displaying the model performance metrics by filtering type. Columns of the matrices represent an increasing level of filtering. MC 0.95+ and 0.99+ corresponds to filtering in all detections with model confidence > 0.95 or 0.99 respectively; HNR 0.1+ corresponds to filtering in all detections with harmonic noise ratio > 0.1; DL 6+ corresponds to filtering in all detections which are at least 6 seconds long. The filtering parameters used on the entire Yellowstone dataset are displayed in the last column (MC 0.99+, HNR 0.1+ and DL 6+).

### 3.2. Effect of snowmobile detection on number of bird vocalizations

Histograms of the average number of bird vocalizations across sites for the three different times of day displayed in Fig. 4 suggest a decrease of the number of bird vocalizations before the detection of a snowmobile in the morning and in the evening (average number of vocalizations in morning = 1.55, 1.58, 1.35; average number of vocalizations in evening = 1.44, 1.39, 1.26 in the ranges [240 s, 225 s], [135 s, 120 s] and [15 s, 0 s]) and an increase after detection of the snowmobile for morning, afternoon and evening (average number of vocalizations in morning = 1.34, 1.49, 1.55; average number of vocalizations in afternoon = 1.31, 1.50, 1.50 and average number of vocalizations in evening = 1.32, 1.43, 1.39 in the ranges [0 s, 15 s], [120 s, 135 s] and [225 s, 240 s]).

**Fig 4:**
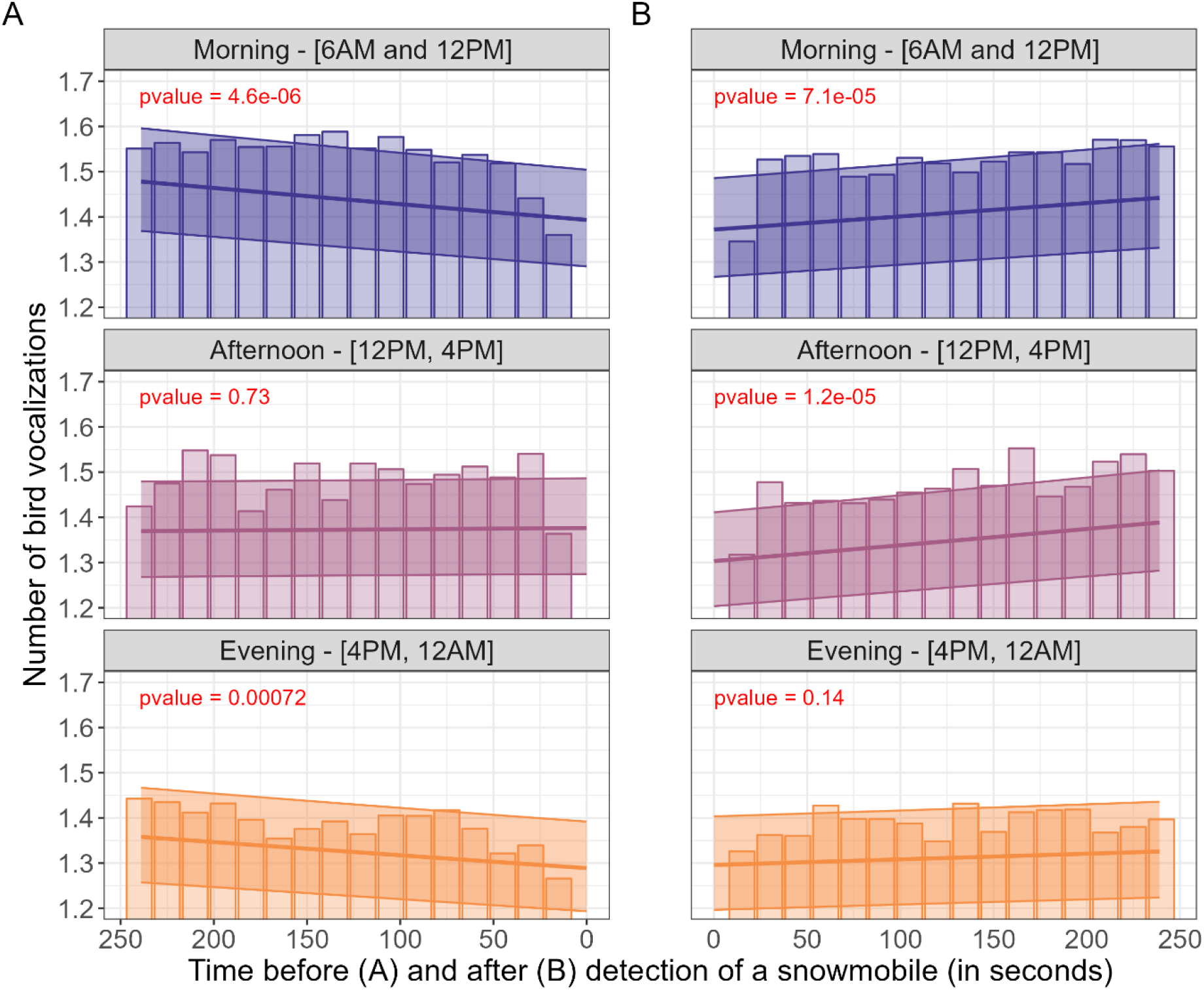
Histograms of the number of bird calls and predicted number of bird calls in 15 seconds intervals before (A) and after (B) detection of a snowmobile, averaged across all the sites in the Yellowstone dataset. The average number of bird vocalizations predicted by our model is displayed by the line. The 95% confidence interval is also displayed. The p-values represent the significance level for the estimated marginal means of the interaction between time of day (t-test with 95% confidence interval).

The statistical models confirm that the passage of snowmobile significatively influence the number of bird vocalizations as there is a significant decrease of the number bird vocalizations before detection of a snowmobile in the morning and evening (i.e. p-values for the estimated marginal means for the interaction between time of day and before detection of a snowmobile < 0.0001, = 0.0007 for morning and evening, Fig 3 and Supplementary Table 1) and a significant increase of the number of bird vocalizations after the detection of a snowmobile in the morning and in the afternoon (i.e. p-values for the estimated marginal means for the interaction between time of day and before detection of a snowmobile = 0.0001, < 0.0001 for morning and afternoon, Fig 3 and Supplementary Table 1).

Model predictions indicate that the predicted number of bird vocalization 240 seconds and 0 second before detection of a snowmobile lead to a decrease of 0.08 vocalizations (1.47 to 1.39 vocalizations per 15 second bin) and 0.16 (1.35 to 1.19 vocalizations) translating in the loss of 5.5% and 12% of of the predicted number of vocalizations in the morning and evening respectively. The predicted number of vocalizations however remains stable in the afternoon 240 seconds and 0 second before the detection of a snowmobile. In contrast, the model predicts that the number of bird vocalizations 0 second and 240 seconds after detection of a snowmobile increase by an average of 0.07 (from 1.37 to 1.44) and 0.08 (from 1.30 to 1.38) vocalizations for morning and afternoon. This translates into a gain of 5% and 6% of vocalizations.

## 4. Discussion

In this study, we used a combination of deep learning and statistical techniques to assess the impact of snowmobile noise on bird vocalizations. Our findings provide valuable insights into the relationships between snowmobile noise and avian vocalization patterns during the non-breeding winter season. The automated snowmobile detection model we developed proved effective in identifying snowmobile noise in the acoustic monitoring data collected in Yellowstone National Park. By applying this model to a large winter acoustic dataset, we were able to analyze the temporal effects of snowmobile noise on bird vocalizations.

Our findings revealed that snowmobile noise had a significant effect on avian vocalizations during the non-breeding winter season, which has been an understudied area in the literature. The passage of snowmobiles led to a decrease in the frequency of bird vocalizations both during mornings and evenings, indicating a disturbance to their communication. Nevertheless, we observe an increase in the frequency of bird vocalization after the detection of a snowmobile, indicating a recovery in avian vocalization and potentially indicating some resilience to the effect of sporadic noise events such as passage of snowmobiles.

Increased noise levels have been shown to have implications for breeding success and habitat selection in birds (Kight & Swaddle, 2011; Ware et al., 2015), but much less is known about noise impacts on wintering birds. Understanding these effects in the non-breeding season is crucial for a comprehensive assessment of the impacts of noise on avian communities throughout their life cycle. Many wintering birds exhibit flocking behavior to improve foraging success and avoid predators (Davies et al., 2012), and within-flock calls help birds avoid predators, find food, and coordinate movements (Freeberg & Lucas, 2002). Noise disturbances to flock communication may thus disrupt important winter behaviors like foraging, predator avoidance and pre-breeding season mate selection. Disturbances to winter flock communication may thus have indirect impacts on breeding season outcomes: breeding pairs often form between birds who are members of the same winter flocks (Firth et al., 2018; Psorakis et al., 2012; Saitou, 1979), and these partnerships can improve breeding success in wild passerine populations (Culina et al., 2020). Though we discovered a statistically significant impact of snowmobile noise on avian vocalization detections, it is unclear whether the observed effect size would constitute a substantial biological impact on distribution and abundance. It was outside the scope of this study to assess other potential impacts of noise on wintering birds, such as avoidance altogether of noisy corridors. However, research suggests that winter avian abundance and distribution is driven more by food availability than by road or rail noise (Wiącek et al., 2019; Wiącek & Polak, 2015), so if food availability happens to be high near a snowmobile corridor, it is possible that wintering birds will risk the potential impacts of snowmobile noise to their communication.

The results of this study underscore the importance of considering noise impacts in the broader context of habitat loss and climate change, as anthropogenic noise can act as a significant stressor, potentially synergistically with other stressors on wildlife populations (Francis et al., 2009). By quantifying the extent of the influence of snowmobile engine noise on avian vocalizations in the non-breeding season, our research contributes to a more comprehensive understanding of the stressors facing avian communities.

This case study may provide insights for natural resource managers who balance snowmobile recreation and avian conservation priorities. The effect size observed in the study is small but statistically significant; results must be contextualized within circumstances of the Yellowstone dataset. In the middle of the acoustic monitoring period covered by this study, Yellowstone National Park adopted a Winter Use Plan (National Park Service, 2013), which included restrictions on the total number of daily oversnow vehicle transportation events allowed in the park, a list of allowable "best available technology" snowmobiles, a 67 dBA limit on snowmobiles operated within the park, and an Adaptive Management Program to assess whether oversnow vehicle use is exceeding the predicted noise budget. Existing management practices, such as event grouping and use of best available technologies, thus may have mitigated the impacts found in this study. Impacts of noise on winter bird vocalizations might be larger in locations with no noise mitigation measures in place. In such areas, managers might choose to minimize disturbance and protect critical habitat areas. For example, implementing speed restrictions or designated snowmobile-free zones in areas with high avian activity during specific hours in the winter season (e.g. morning) may reduce disturbance to bird vocalization activity.

It should be noted that the time window used in this analysis (i.e. [240s,0s], [0s,240s] before and after detection of a snowmobile respectively) has been chosen based on extrapolations from human auditory capabilities. We acknowledge the differences in hearing capacities and perception between humans and birds (Dooling, 2002), as well as the wide range of variables that can influence such estimates (e.g. wind, landscape profile). However, lacking more specific data, we believe we offer a reasonable starting point for analyzing the impact of snowmobile noise on bird communities. The methodology we developed for assessing noise impacts from snowmobiles using deep learning techniques offers significant advantages for both scientific research and management purposes. By automating the detection and classification of snowmobile noise, we were able to analyze large amounts of acoustic monitoring data efficiently and accurately. This automated approach not only saves time and effort compared to manual sound annotation but also allows for the examination of a larger temporal and geographic scope, providing a more comprehensive understanding of the impact of snowmobile noise on avian vocalizations. Nevertheless, the bird classifier model we used (i.e. BirdNET) is unable to identify the type of call emitted and it is not possible to differentiate between alarm, call or song. Such analysis would require custom built deep learning models trained on the specific calls of each bird species. We have not investigated the effect of snowmobile noise on particular species either as we aggregated the number of calls independently of the species. This is due to the lack of documentation on BirdNET’s performance on particular species and results of such analysis could be misleading. While it is possible to obtain BirdNET’s precision and accuracy for different species (Sethi et al., 2021), it is resource consuming and can hardly be scaled for large numbers of bird species.

In conclusion, our study demonstrates a small impact of snowmobile noise on avian vocalizations during the non-breeding winter season in Yellowstone National Park. The application of deep learning techniques allowed for efficient detection and analysis of snowmobile noise, providing valuable insights into the relationships between anthropogenic noise and avian vocalization patterns. This research has important implications for the management of natural areas where recreation and wildlife conservation coexist, highlighting the need for measures to mitigate noise impacts on avian communities and protect critical habitats.

## Authors contribution

BC, FG and TAR developed the snowmobile detection model. JW labeled the training and test dataset. BC, DPJ and IAB analyzed the detections resulting from the snowmobile and BirdNET models. The Yellowstone dataset was provided by CB, DB and BB. The manuscript was drafted by BC and IAB. All authors provided feedback and approved the manuscript.

## Acknowledgments

We thank Francesco Frassinelli who gave support in running the training pipeline on High Performance Clusters (HPCs). This work was supported by the Norwegian Research Council (grant number 160022/F40). S.S.S. was supported by a Herchel Smith Fellowship from the School of Biological Sciences at the University of Cambridge. Thank you to all staff at Yellowstone National Park who collected the passive acoustic monitoring data that enabled this case study.

## Conflict of Interest

The authors declare no conflict of interest.

## Data availability statement

Code for running the snowmobile model is available on GitHub (https://github.com/NINAnor/snowmobile_analyzer). The repository also contains the R scripts and dataset used to run the statistical modeling.

## Notes

https://github.com/NINAnor/snowmobile_analyzer

### Competing Interest Statement

The authors have declared no competing interest.

