## Supplementaty material for "Snowmobile noise alters bird vocalization patterns during winter and pre-breeding season"

**Supplementary material for:** Snowmobile noise alters bird vocalizations patterns during winter and pre-breeding season

Page 1: **Supplementary map 1.** Map of the sites analyzed by both the snowmobile and bird classification model.

Page 2: **Supplementary Table 1**. Estimated marginal means and contrasts for the interaction between time of day and snowmobile detection. P-values have been determined using t-tests (95% confidence interval)

Page 3: **Supplementary Table 2**. Results from the linear models. In bold text are the significant coefficients (p-value < 0.05).

Page 4: **Supporting document 1:** Table listing the species list used as input to BirdNET.

**Supplementary map 1.** Map of the sites analyzed by both the snowmobile and bird classification model.

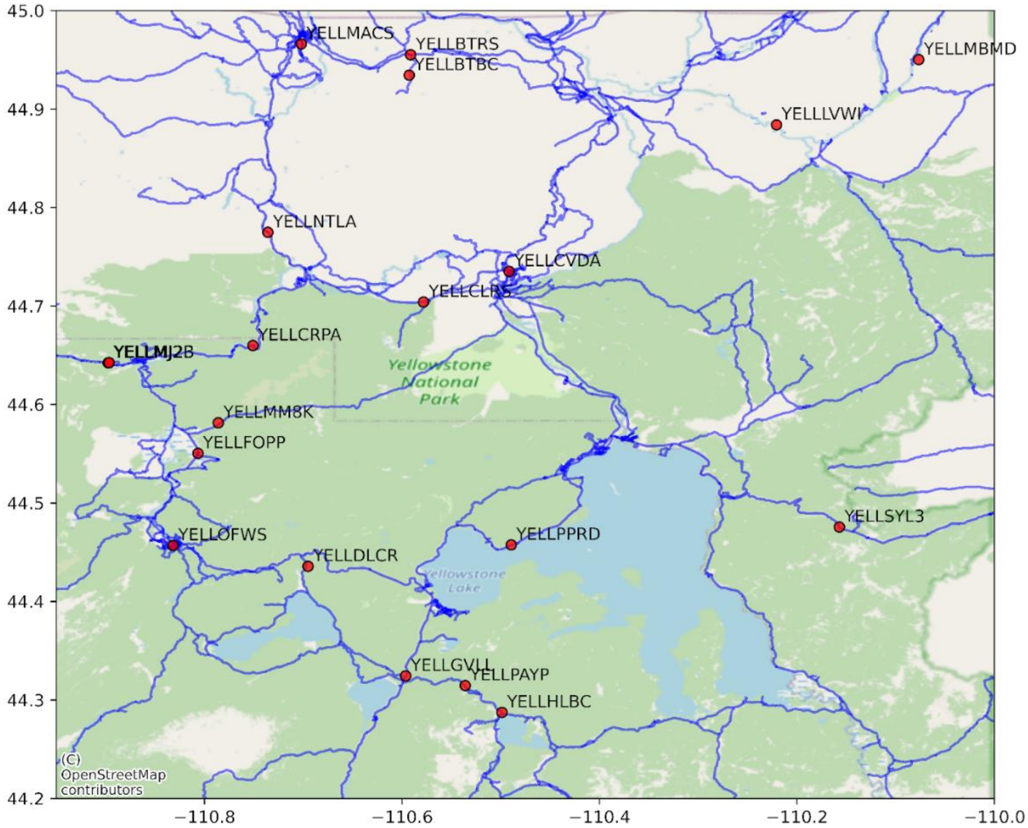

**Supplementary Table 1:** Estimated marginal means and contrasts for the interaction between time of day and snowmobile detection. P-values have been determined using t-tests (95% confidence interval)

| Model | Time of day | Estimate | SE | df | t.ratio | p.value |
|---|---|---|---|---|---|---|
| Before passage | | | | | | |
| | **Morning** | **0.000247** | **5.40e-05** | **34901** | **4.582** | **<.0001** |
| | Afternoon | -0.000021 | 6.08e-05 | 34901 | -0.345 | 0.7304 |
| | **Evening** | **0.000219** | **6.46e-05** | **34901** | **3.382** | **0.0007** |
| After passage | | | | | | |
| | **Morning** | **2.08e-04** | **5.25e-05** | **34929** | **3.972** | **0.0001** |
| | **Afternoon** | **2.66e-04** | **6.08e-05** | **34929** | **4.372** | **<.0001** |
| | Evening | 9.52e-05 | 6.52e-05 | 34929 | 1.459 | 0.1445 |

| Model | Contrasts | estimate | SE | df | t.ratio | p.value |
|---|---|---|---|---|---|---|
| Before passage | | | | | | |
| | **Morning-Afternoon** | **2.68e-04** | **8.12e-05** | **34901** | **3.307** | **0.0027** |
| | Morning-Evening | 2.87e-05 | 8.41e-05 | 34901 | 0.342 | 0.9377 |
| | **Afternoon-Evening** | **-2.40e-04** | **8.86e-05** | **34901** | **-2.704** | **0.0188** |
| After passage | | | | | | |
| | Morning-Afternoon | -5.74e-05 | 8.01e-05 | 34929 | -0.716 | 0.7539 |
| | Morning-Evening | 1.13e-04 | 8.36e-05 | 34929 | 1.354 | 0.3651 |
| | Afternoon-Evening | 1.71e-04 | 8.90e-05 | 34929 | 1.916 | 0.1340 |

**Supplementary Table 2**. Results from the linear models. In bold text are the significant coefficients (p-value < 0.05).

| Model | Variable | Estimate | Std. | Error | Pr(>\|z\|) |
|---|---|---|---|---|---|
| Before passage | | | | | |
| | **(Intercept)** | **0.297178** | **0.023348** | **12.728** | **<2e-16*****|
| | **scale(closest_ss_before)** | **0.016813** | **0.003670** | **4.581** | **4.62e-06***** |
| | **time_of_dayMM** | **-0.040968** | **0.006906** | **-5.932** | **2.98e-09***** |
| | **time_of_dayPM** | **-0.080877** | **0.006342** | **-12.753** | **<2e-16***** |
| | scale(sknt) | -0.003666 | 0.003272 | -1.120 | 0.262536 |
| | **scale(tmpf)** | **0.012756** | **0.004053** | **3.148** | **0.001645**** |
| | scale(relh) | 0.003237 | 0.003862 | 0.838 | 0.401898 |
| | **scale(closest_ss_before):time_of_dayMM** | **-0.018237** | **0.005515** | **-3.307** | **0.000944***** |
| | scale(closest_ss_before):time_of_dayPM | -0.001952 | 0.005718 | -0.341 | 0.732796 |
| After passage | | | | | |
| | **(Intercept)** | **0.2856038** | **0.0210905** | **13.542** | **<2e-16***** |
| | **scale(closest_ss_after)** | **0.0141315** | **0.0035572** | **3.973** | **7.11e-05***** |
| | **time_of_dayMM** | **-0.0451157** | **0.0068554** | **-6.581** | **4.67e-11***** |
| | **time_of_dayPM** | **-0.0693716** | **0.0063077** | **-10.998** | **<2e-16***** |
| | scale(sknt) | -0.0009682 | 0.0032618 | -0.297 | 0.76661 |
| | **scale(tmpf)** | **0.0131781** | **0.0040736** | **3.235** | **0.00122**** |
| | scale(relh) | 0.0051406 | 0.0038453 | 1.337 | 0.18127 |
| | scale(closest_ss_after):time_of_dayMM | 0.0038885 | 0.0054320 | 0.716 | 0.47409 |
| | scale(closest_ss_after):time_of_dayPM | -0.0076812 | 0.0056689 | -1.355 | 0.17543 |

**Supporting document 1.** Table listing the species list used as input to BirdNET.

| Scientific name | Common name |
|---|---|
| Acanthis flammea | Common Redpoll |
| Accipiter cooperii | Cooper's Hawk |
| Accipiter gentilis | Northern Goshawk |
| Accipiter striatus | Sharp-shinned Hawk |
| Actitis macularius | Spotted Sandpiper |
| Aechmophorus clarkii | Clark's Grebe |
| Aechmophorus occidentalis | Western Grebe |
| Aegolius acadicus | Northern Saw-whet Owl |
| Aegolius funereus | Boreal Owl |
| Aeronautes saxatalis | White-throated Swift |
| Agelaius phoeniceus | Red-winged Blackbird |
| Aix sponsa | Wood Duck |
| Alectoris chukar | Chukar |
| Ammodramus savannarum | Grasshopper Sparrow |
| Amphispiza bilineata | Black-throated Sparrow |
| Anas acuta | Northern Pintail |
| Anas crecca | Green-winged Teal |
| Anas platyrhynchos | Mallard |
| Anser albifrons | Greater White-fronted Goose |
| Anthus rubescens | American Pipit |
| Antigone canadensis | Sandhill Crane |
| Aquila chrysaetos | Golden Eagle |
| Ardea alba | Great Egret |
| Ardea herodias | Great Blue Heron |
| Arenaria interpres | Ruddy Turnstone |
| Artemisiospiza nevadensis | Sagebrush Sparrow |
| Asio flammeus | Short-eared Owl |
| Asio otus | Long-eared Owl |
| Athene cunicularia | Burrowing Owl |
| Aythya affinis | Lesser Scaup |
| Aythya collaris | Ring-necked Duck |
| Aythya marila | Greater Scaup |
| Aythya valisineria | Canvasback |
| Bartramia longicauda | Upland Sandpiper |
| Bombycilla cedrorum | Cedar Waxwing |
| Bombycilla garrulus | Bohemian Waxwing |
| Bonasa umbellus | Ruffed Grouse |
| Botaurus lentiginosus | American Bittern |
| Branta canadensis | Canada Goose |
| Branta hutchinsii | Cackling Goose |
| Bubo virginianus | Great Horned Owl |
| Bubulcus ibis | Cattle Egret |

| | |
|---|---|
| Bucephala albeola | Bufflehead |
| Bucephala clangula | Common Goldeneye |
| Buteo jamaicensis | Red-tailed Hawk |
| Buteo lineatus | Red-shouldered Hawk |
| Buteo platypterus | Broad-winged Hawk |
| Buteo swainsoni | Swainson's Hawk |
| Butorides virescens | Green Heron |
| Calamospiza melanocorys | Lark Bunting |
| Calcarius lapponicus | Lapland Longspur |
| Calidris alba | Sanderling |
| Calidris bairdii | Baird's Sandpiper |
| Calidris fuscicollis | White-rumped Sandpiper |
| Calidris mauri | Western Sandpiper |
| Calidris melanotos | Pectoral Sandpiper |
| Calidris minutilla | Least Sandpiper |
| Calidris pusilla | Semipalmated Sandpiper |
| Caracara plancus | Crested Caracara |
| Cardellina pusilla | Wilson's Warbler |
| Cardellina pusilla | Wilson's Warbler |
| Catharus fuscescens | Veery |
| Catharus guttatus | Hermit Thrush |
| Catharus ustulatus | Swainson's Thrush |
| Catherpes mexicanus | Canyon Wren |
| Certhia americana | Brown Creeper |
| Charadrius semipalmatus | Semipalmated Plover |
| Charadrius vociferus | Killdeer |
| Chlidonias niger | Black Tern |
| Chondestes grammacus | Lark Sparrow |
| Chordeiles minor | Common Nighthawk |
| Chroicocephalus philadelphia | Bonaparte's Gull |
| Cinclus mexicanus | American Dipper |
| Circus hudsonius | Northern Harrier |
| Cistothorus palustris | Marsh Wren |
| Clangula hyemalis | Long-tailed Duck |
| Coccothraustes vespertinus | Evening Grosbeak |
| Coccyzus erythropthalmus | Black-billed Cuckoo |
| Colaptes auratus | Northern Flicker |
| Columba livia | Rock Pigeon |
| Contopus cooperi | Olive-sided Flycatcher |
| Contopus sordidulus | Western Wood-Pewee |
| Corthylio calendula | Ruby-crowned Kinglet |
| Corvus brachyrhynchos | American Crow |
| Corvus corax | Common Raven |
| Coturnicops noveboracensis | Yellow Rail |
| Cyanocitta cristata | Blue Jay |
| Cyanocitta stelleri | Steller's Jay |
| Cygnus buccinator | Trumpeter Swan |

| | |
|---|---|
| Cygnus columbianus | Tundra Swan |
| Cygnus cygnus | Whooper Swan |
| Dendragapus obscurus | Dusky Grouse |
| Dolichonyx oryzivorus | Bobolink |
| Dryobates pubescens | Downy Woodpecker |
| Dryobates villosus | Hairy Woodpecker |
| Dryocopus pileatus | Pileated Woodpecker |
| Dumetella carolinensis | Gray Catbird |
| Egretta thula | Snowy Egret |
| Egretta tricolor | Tricolored Heron |
| Empidonax hammondii | Hammond's Flycatcher |
| Empidonax minimus | Least Flycatcher |
| Empidonax oberholseri | Dusky Flycatcher |
| Empidonax occidentalis | Cordilleran Flycatcher |
| Empidonax traillii | Willow Flycatcher |
| Eremophila alpestris | Horned Lark |
| Euphagus carolinus | Rusty Blackbird |
| Euphagus cyanocephalus | Brewer's Blackbird |
| Falco columbarius | Merlin |
| Falco peregrinus | Peregrine Falcon |
| Falco sparverius | American Kestrel |
| Fulica americana | American Coot |
| Gallinago delicata | Wilson's Snipe |
| Gavia immer | Common Loon |
| Geothlypis tolmiei | MacGillivray's Warbler |
| Geothlypis trichas | Common Yellowthroat |
| Glaucidium gnoma | Northern Pygmy-Owl |
| Gymnorhinus cyanocephalus | Pinyon Jay |
| Haemorhous cassinii | Cassin's Finch |
| Haemorhous mexicanus | House Finch |
| Haliaeetus leucocephalus | Bald Eagle |
| Himantopus mexicanus | Black-necked Stilt |
| Hirundo rustica | Barn Swallow |
| Hydroprogne caspia | Caspian Tern |
| Hydroprogne caspia | Caspian Tern |
| Icteria virens | Yellow-breasted Chat |
| Icterus bullockii | Bullock's Oriole |
| Ixoreus naevius | Varied Thrush |
| Junco hyemalis | Dark-eyed Junco |
| Lanius excubitor | Great Gray Shrike |
| Lanius ludovicianus | Loggerhead Shrike |
| Larus argentatus | Herring Gull |
| Larus californicus | California Gull |
| Larus canus | Common Gull |
| Larus delawarensis | Ring-billed Gull |
| Leiothlypis peregrina | Tennessee Warbler |
| Leiothlypis ruficapilla | Nashville Warbler |

| | |
|---|---|
| Leucophaeus pipixcan | Franklin's Gull |
| Leucosticte tephrocotis | Gray-crowned Rosy-Finch |
| Limnodromus griseus | Short-billed Dowitcher |
| Limnodromus scolopaceus | Long-billed Dowitcher |
| Limosa fedoa | Marbled Godwit |
| Limosa haemastica | Hudsonian Godwit |
| Lophodytes cucullatus | Hooded Merganser |
| Loxia curvirostra | Red Crossbill |
| Loxia leucoptera | White-winged Crossbill |
| Mareca americana | American Wigeon |
| Mareca penelope | Eurasian Wigeon |
| Mareca strepera | Gadwall |
| Megaceryle alcyon | Belted Kingfisher |
| Megascops kennicottii | Western Screech-Owl |
| Melanerpes erythrocephalus | Red-headed Woodpecker |
| Melanerpes lewis | Lewis's Woodpecker |
| Meleagris gallopavo | Wild Turkey |
| Melospiza georgiana | Swamp Sparrow |
| Melospiza lincolnii | Lincoln's Sparrow |
| Melospiza melodia | Song Sparrow |
| Mergus merganser | Common Merganser |
| Mergus serrator | Red-breasted Merganser |
| Mniotilta varia | Black-and-white Warbler |
| Molothrus ater | Brown-headed Cowbird |
| Myadestes townsendi | Townsend's Solitaire |
| Myiarchus cinerascens | Ash-throated Flycatcher |
| Nannopterum auritum | Double-crested Cormorant |
| Nucifraga columbiana | Clark's Nutcracker |
| Numenius americanus | Long-billed Curlew |
| Nycticorax nycticorax | Black-crowned Night-Heron |
| Oreoscoptes montanus | Sage Thrasher |
| Oxyura jamaicensis | Ruddy Duck |
| Pandion haliaetus | Osprey |
| Parkesia noveboracensis | Northern Waterthrush |
| Passer domesticus | House Sparrow |
| Passerculus sandwichensis | Savannah Sparrow |
| Passerella iliaca | Fox Sparrow |
| Passerina amoena | Lazuli Bunting |
| Patagioenas fasciata | Band-tailed Pigeon |
| Perdix perdix | Gray Partridge |
| Perisoreus canadensis | Canada Jay |
| Petrochelidon pyrrhonota | Cliff Swallow |
| Phalaropus lobatus | Red-necked Phalarope |
| Phalaropus tricolor | Wilson's Phalarope |
| Pheucticus ludovicianus | Rose-breasted Grosbeak |
| Pheucticus melanocephalus | Black-headed Grosbeak |
| Pica hudsonia | Black-billed Magpie |

| | |
|---|---|
| Picoides arcticus | Black-backed Woodpecker |
| Picoides dorsalis | American Three-toed Woodpecker |
| Pinicola enucleator | Pine Grosbeak |
| Pipilo chlorurus | Green-tailed Towhee |
| Pipilo maculatus | Spotted Towhee |
| Piranga ludoviciana | Western Tanager |
| Plectrophenax nivalis | Snow Bunting |
| Plegadis chihi | White-faced Ibis |
| Pluvialis squatarola | Black-bellied Plover |
| Podiceps auritus | Horned Grebe |
| Podiceps grisegena | Red-necked Grebe |
| Podiceps nigricollis | Eared Grebe |
| Podilymbus podiceps | Pied-billed Grebe |
| Poecile atricapillus | Black-capped Chickadee |
| Poecile gambeli | Mountain Chickadee |
| Polioptila caerulea | Blue-gray Gnatcatcher |
| Pooecetes gramineus | Vesper Sparrow |
| Porzana carolina | Sora |
| Protonotaria citrea | Prothonotary Warbler |
| Psiloscops flammeolus | Flammulated Owl |
| Quiscalus quiscula | Common Grackle |
| Rallus limicola | Virginia Rail |
| Recurvirostra americana | American Avocet |
| Regulus satrapa | Golden-crowned Kinglet |
| Riparia riparia | Bank Swallow |
| Rissa tridactyla | Black-legged Kittiwake |
| Salpinctes obsoletus | Rock Wren |
| Sayornis saya | Say's Phoebe |
| Seiurus aurocapilla | Ovenbird |
| Selasphorus calliope | Calliope Hummingbird |
| Selasphorus platycercus | Broad-tailed Hummingbird |
| Selasphorus rufus | Rufous Hummingbird |
| Setophaga citrina | Hooded Warbler |
| Setophaga coronata | Yellow-rumped Warbler |
| Setophaga discolor | Prairie Warbler |
| Setophaga fusca | Blackburnian Warbler |
| Setophaga pensylvanica | Chestnut-sided Warbler |
| Setophaga petechia | Yellow Warbler |
| Setophaga ruticilla | American Redstart |
| Setophaga striata | Blackpoll Warbler |
| Setophaga tigrina | Cape May Warbler |
| Setophaga townsendi | Townsend's Warbler |
| Sialia currucoides | Mountain Bluebird |
| Sialia mexicana | Western Bluebird |
| Sitta canadensis | Red-breasted Nuthatch |
| Sitta carolinensis | White-breasted Nuthatch |
| Sitta pygmaea | Pygmy Nuthatch |

| | |
|---|---|
| Spatula clypeata | Northern Shoveler |
| Spatula cyanoptera | Cinnamon Teal |
| Spatula discors | Blue-winged Teal |
| Sphyrapicus nuchalis | Red-naped Sapsucker |
| Sphyrapicus thyroideus | Williamson's Sapsucker |
| Spinus pinus | Pine Siskin |
| Spinus psaltria | Lesser Goldfinch |
| Spinus tristis | American Goldfinch |
| Spizella breweri | Brewer's Sparrow |
| Spizella pallida | Clay-colored Sparrow |
| Spizella passerina | Chipping Sparrow |
| Spizelloides arborea | American Tree Sparrow |
| Stelgidopteryx serripennis | Northern Rough-winged Swallow |
| Sterna forsteri | Forster's Tern |
| Sterna hirundo | Common Tern |
| Sterna paradisaea | Arctic Tern |
| Streptopelia decaocto | Eurasian Collared-Dove |
| Streptopelia decaocto | Eurasian Collared-Dove |
| Strix nebulosa | Great Gray Owl |
| Sturnella neglecta | Western Meadowlark |
| Sturnus vulgaris | European Starling |
| Tachycineta bicolor | Tree Swallow |
| Tachycineta thalassina | Violet-green Swallow |
| Tringa flavipes | Lesser Yellowlegs |
| Tringa melanoleuca | Greater Yellowlegs |
| Tringa semipalmata | Willet |
| Tringa solitaria | Solitary Sandpiper |
| Troglodytes aedon | House Wren |
| Turdus migratorius | American Robin |
| Tyrannus forficatus | Scissor-tailed Flycatcher |
| Tyrannus tyrannus | Eastern Kingbird |
| Tyrannus verticalis | Western Kingbird |
| Vireo gilvus | Warbling Vireo |
| Vireo olivaceus | Red-eyed Vireo |
| Xanthocephalus xanthocephalus | Yellow-headed Blackbird |
| Zenaida macroura | Mourning Dove |
| Zonotrichia albicollis | White-throated Sparrow |
| Zonotrichia leucophrys | White-crowned Sparrow |
| Zonotrichia querula | Harris's Sparrow |
